# Geodesic-based distance reveals non-linear topological features in neural activity from mouse visual cortex

**DOI:** 10.1101/2021.05.21.444993

**Authors:** Kosio Beshkov, Paul Tiesinga

## Abstract

An increasingly popular approach to the analysis of neural data is to treat activity patterns as being constrained to and sampled from a manifold, which can be characterized by its topology. The *persistent homology* method identifies the type and number of holes in the manifold thereby yielding functional information about the coding and dynamic properties of the underlying neural network. In this work we give examples of highly non-linear manifolds in which the *persistent homology* algorithm fails when it uses the Euclidean distance which does not always yield a good approximation of the true distance distribution of a point cloud sampled from a manifold. To deal with this issue we propose a simple strategy for the estimation of the geodesic distance which is a better approximation of the true distance distribution and can be used to successfully identify highly non-linear features with *persistent homology*. To document the utility of our method we model a circular manifold, based on orthogonal sinusoidal basis functions and compare how the chosen metric determines the performance of the *persistent homology* algorithm. Furthermore we discuss the robustness of our method across different manifold properties and point out strategies for interpreting its results as well as some possible pitfalls of its application. Finally we apply this analysis to neural data coming from the *Visual Coding - Neuropixels* dataset recorded in mouse visual cortex after stimulation with drifting gratings at the Allen Institute. We find that different manifolds with a non-trivial topology can be seen across regions and stimulus properties. Finally, we discuss what these manifolds say about visual computation and how they depend on stimulus parameters.

## 1 Introduction

While in the past neuroscientists were limited to recordings of only a few cells, today because of advancements in large scale neural recording methods and open data, datasets of large populations of simultaneously recorded cells are publicly available for further analysis. This motivates the development of new analysis methods with which to advance our mechanistic understanding of neural processing. An intuitive way to think about the activity of a large population of neurons is as a set of points in high dimensional space which form a manifold. Determining relevant structure in data distributed across a manifold generated by a neural network can contribute to our understanding of neural computation, by relating encoded variables to positions on the manifold. In other words, points on a manifold correspond to particular network states, which in turn are said to represent a value of an external or internal variable. A popular way of doing this, which has yielded several interesting results showing the relation between manifold structure and neural representation, is through the application of both linear and non-linear dimensionality reduction methods (Gao and Ganguli (2015), Gallego et al. (2017), Chaudhuri et al. (2019), Saxena and Cunningham (2019), Mastrogiuseppe and Ostojic (2018)). One problem with such methods is that they rely on the point cloud being sampled densely enough that one can find an embedding in an appropriate number of dimensions (usually not more than 3 if visualization is desirable) while preserving its relevant topological properties. Typical topological properties are the number and type of holes, which under the assumption that neural manifolds are smooth, are expected to be representationally invariant, (Shepard and Chipman (1970), Kriegeskorte et al. (2008)) and a robust characterization of neural activity. Another more recent technique coming from the field of *topological data analysis* is *persistent homology* (Ghrist (2008), Carlsson (2009)), Wasserman (2018)), which finds the different types of holes in a point cloud. This method has been applied to study the topological structure of neural activity in the visual cortex (Singh et al. (2008)), place cells in the hippocampus (Curto and Itskov (2008), Dabaghian et al. (2012), Babichev et al. (2016), Babichev and Dabaghian (2018)) and head direction cells (Chaudhuri et al. (2019)). All of these studies indicate that the topology of neural activity is related to that of the presented stimuli. There are two types of point clouds that one can construct from neural data. In the first case the time trace of each neuron is taken to be a point and the traced-out manifold represents the relations between the neurons, in this framing the axes are the time bins and the number of points in the space is the number of neurons. This is useful when the receptive fields of neurons cover an input space with a particularly interesting topology (Curto (2017)). In the second type of point cloud, each time bin across the population of neurons is taken to be a point, here the neurons are the axes and the number of points is the same as the number of bins. As a result, the manifold represents the relations between network states. In this study we use the second approach. When applying *persistent homology*, one creates a set of simplicial complexes, which are generalizations of graphs that capture higher-order relations, based on the Euclidean distance between points in a point cloud, where all points within a distance below a certain threshold are considered to be connected. In this way, different simplicial complexes are generated and different topological features appear as a function of the threshold. Depending on how long a topological feature persists for varying thresholds, one can assess whether it is a true feature of the point cloud or more properly considered as topological noise.

This approach assumes that the Euclidean distance is a good enough approximation of the real distance between points. We argue that this assumption can fail in highly nonlinear manifolds by not finding the structure which is there or by finding false positive results, because the Euclidean distance creates artificial shortcuts between points in different regions of the manifold. As a result it might also fail in correctly describing the topological relations that a neural network or its dynamics encode. To fix this problem we propose to use a numerical approximation of the geodesic distance with which the *persistent homology* algorithm can correctly identify structure in high dimensional non-linear manifolds. We quantify, using model generated data, how our algorithm depends on several manifold properties like sampling density, distribution width, curvature and noise. We also discuss the potential limitations and concomitant solutions of our approach, like the necessity to chose specific parameter values for each analysis and apply post-hoc significance testing. Lastly we apply our modified method to explore the topological properties found in the mouse visual system under stimulation with drifting grating stimuli. To do this we used the Allen Institute Visual Coding - Neuropixels dataset (Siegle et al. (2021)), which provides an immense amount of high-quality neural recordings.

## 2 Methods

### 2.1 Simplicial Complexes and Homology

A simplicial complex is a generalization of a graph. A graph can be defined by a vertex and edge set {*V, E*}, where the vertices denote the single objects and the edges denote binary relations. The generalization is straightforward, one can add sets which describe higher order relations. For example vertices are 0-simplices, edges are 1-simplices, triangles are 2-simplices and in general the relation between k points is called a (k-1)-simplex. One can consider a (k-1)-simplex as a vector [*v*_1_, *v*_2_, .., *v*_*k*_] and the set of all (k-1) simplices as a vector space over a field of coefficients *G*(this is known more generally as a chain complex *C*_*k*_). This leaves us with a set of vector spaces which are related by the boundary map *∂*_*k*_ : *C*_*k*_ → *C*_*k−*1_. This map has the purpose of sending closed cycles to the null space (kernel) of the lower dimensional vector space and can be explicitly written as

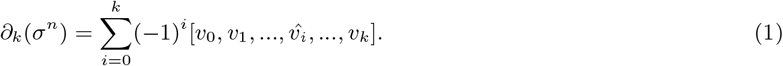

The hat operator deletes the element it is over.

Homology is then calculated by asking how many closed cycles there are which are not the boundary of a higher dimensional object (another way to say this is that the closed cycles are not filled in by a higher dimensional object). The formal expression that implements this calculation is:

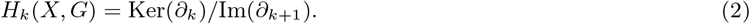

### 2.2 Persistent Homology

The *persistent homology* algorithm takes a distance matrix and varies a distance treshold to obtain a set of simplicial complexes. The distance matrices represent a set of points in a metric space, from which the Vietoris-Rips complex (Zomorodian (2010)), is calculated by choosing a neighborhood around each such point and adding a (k-1)-simplex whenever the distance between k points is less or equal to the diameter of the neighborhood. Increasing the radius induces a sequence of simplicial complexes:

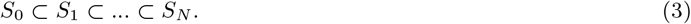

*Persistent homology* then calculates the *homology* groups for each such simplicial complex, yielding a set of homology groups describing the topological features for each parameter value:

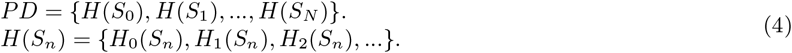

Here *H*(*S*_*n*_) denotes a set of homology groups corresponding to holes of different dimensions and PD is an abbreviation for *persistence diagram*. Geometrically intuitive examples of topological features are the number of connected components (also known as the 0th homology group *H*_0_(*S*)), the number of holes (*H*_1_(*S*)) and the number of cavities like those inside a basketball (*H*_2_(*S*)). In general one can calculate holes of any positive integer dimension, but we will only focus on the aforementioned three. For a more in depth introduction in homology suited for neuroscientists see (Sizemore et al. (2019)) and for a more mathematical treatment see (Hatcher (2002), Edelsbrunner and Harer (2010), Munkres (2018)). Topological features appear in one of the simplicial complexes and disappear in a later one, their persistence is defined as the parameter threshold at which the feature disappears (the death) minus the time at which it first appears (the birth). For any parameter value one can see how many topological features of a given kind exist, this value is known as the Betti number and it is defined as the rank of the homology group (*β*_*n*_ = rank *H*_*n*_(*X*)). For the calculation of the *persistent (co)homology* we used the Python package **Ripser** (Tralie et al. (2018)). The manifolds we looked for were connected spaces without holes (characterized by the sequence of Betti numbers (1,0,0)), circles (1,1,0), spheres (1,0,1), tori (1,2,1) and other (anything with a different Betti number sequence).

### 2.3 Numerical calculation of the geodesic distance

Given a point cloud, we calculate the geodesic distance between its points by first constructing a connected graph and then running a shortest path algorithm to find the shortest distance between each pair of points on the graph. Our implementation uses the **shortest path** scipy function. For the construction of the graph one needs to make a choice regarding the distance at which to create edges between vertices in the point cloud. To construct the graph we placed (D-1)-sphere (D is the dimension of the point cloud embedding) neighborhoods with radius *R* around each point and added an edge between two points whenever they are at a distance *d*(*x, y*) *< R* from each other. To find the minimal scale at which a manifold becomes connected and thus makes the distance between any two points well defined, we took advantage of the fact that the 0-dimensional components in *persistent homology* count the number of connected components. Using this fact we chose the radius by computing the value at which the *persistent homology* algorithm identified only one connected component, in other words we choose the radius at which min_*R*_{*β*_0_[*H*(*S*_*R*_)] = 1}. After that one can choose by how much more to increase the radius, depending on the type of assumptions one makes about the point cloud distribution and the structure one is interested in finding. A higher radius implies the belief that the manifold from which the data is sampled is thicker, while a lower radius implies the opposite.

### 2.4 Generation of high dimensional non-linear manifolds

We define a manifold to be high dimensional, when *principal component analysis* (PCA) (Tipping and Bishop (1999)) requires several dimensions (d*>*3) to explain at least 80% of the variance. In the most extreme case the covariance matrix is equivalent to the identity matrix and in that case each PCA component will explain the same amount of variance. In order to generate high dimensional non-linear manifolds which contain interesting topological information, we used the following orthogonal basis functions:

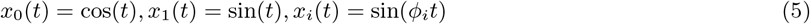

Where *ϕ*_*i*_ ∈ ℤ are integer frequency variables, in order to make the manifold closed. That happens when *t* has gone through 2*π* and the traced out path returns to the starting point. These basis functions are orthogonal and generate a distorted circle in high dimensional space. The orthogonality of the functions is necessary to make the covariance matrix proportional to the identity and is also reminiscent of the trajectories traced by a neural population implementing an efficient code (Stringer et al. (2019))

### 2.5 Comparison between the Euclidean and geodesic metric as a function of manifold parameters

To compare the performance of the two metrics we explored how the difference between the estimated distances changes as a function of the number of samples, curvature, neighborhood radius and noise. We also used persistence images (Adams et al. (2017)) to compare how the estimated homology from the two metrics changes as a function of the aforementioned parameters.This method works by treating the birth-death diagram as an image and applying gaussian kernels (we used a pixel size of 0.1 and a kernel width of 0.05) to each point on the diagram, as a result one gets a value *I*_*x,y*_ for each pixel of the image. To keep the comparisons consistent we normalized the persistence images *I*_*x,y*_ as 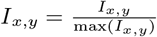. Both the distance matrix and the persistence images are matrices so to calculate the difference between them we used the Frobenius norm on their elementwise difference:

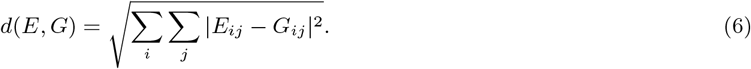

### 2.6 Data preprocessing and analysis

All data analysis was performed on the Allen Institute Visual Coding - Neuropixels dataset (Siegle et al. (2021)). Cells were selected based on their signal to noise ratio (SNR *>* 2) and firing rate (*>* 1Hz). This choice was made so that we only work with cells which are active and reliable. We analysed the neural activity in 5 visual regions (lateral posterior nucleus - LP, primary visual cortex - VISp, rostrolateral area of the visual cortex - VISrl, lateromedial area of the visual cortex - VISl and anteromedial area of the visual cortex - VISam) of all available mice, obtained in response to drifting gratings.

We use the spike count for all 40 presented stimuli (8 directions and 5 temporal frequencies) and all 15 trials (each trial has a duration of 2s) available for each stimulus. Because we want to find the manifold structure generated by the entire stimulus set, we concatenate the responses to all 40 stimuli for each mouse and region separately. Once we have a *neurons*×*[trials*×*stimuli]* (*N* × *T*) matrix for each region, we end up with a point cloud per region per subject representing its activity manifold. Since not all mice had recordings for the full set of regions, we ended up with 126 point clouds. When analysing the fixed temporal frequency manifolds we restrict the stimulus set to 8 stimuli of the same temporal frequency.

### 2.7 Significance testing

To determine whether the topological features we find are significant we perform a permutation-based significance test. Given a point cloud represented by a *N* × *T* matrix, we permute the entries of the matrix across both dimensions 2500 times and run our analysis on the permuted data. We can then calculate the persistence of the topological features. Because permutations enlarge the scale of the persistence diagram, we normalize it beforehand by the maximum persistence in the diagram (we will refer to the normalized persistence of a feature as its strength). The maximal persistence values of the permuted data form a distribution from which one can define a one tailed p value by calculating the corresponding percentile. We use this non-parametric statistical test to see how many of all the datasets have significant topological features, as well as to determine which topological features are significant within each dataset. When calculating the significant topological features in all datasets, we use a false discovery rate multiple test correction (Benjamini and Hochberg (1995)) by taking all of the extracted p-values and applying the **fdrcorrection** function from the **statsmodels** package (Seabold and Perktold (2010)). Finally we also performed a significance test to check if there are differences between the manifolds produced by the activity in each region. To do this we generate a distribution for each topological feature by resampling the sessions and then shuffling the features across regions 1000 times. Since there was an unequal amount of datasets for each region, we took the ratio of appearance of each type of manifold within a region. This way we get a distribution for each topological feature and by computing the percentile of the observed feature score we can define a two-sided p-value with which we can check whether a particular region has a significantly higher or lower amount of topological features of a particular type. The temporal frequency fixed features were compared to the full temporal frequency topological feature distributions. These p-values were corrected with a two-sided false-discovery rate, in which p-values above 0.5 are transformed as *p*^*l*^ = (1 − *p*) and then corrected along with all the other p-values. We used a significance threshold of 0.05, except in the two-sided case where the threshold was set to 0.025.

### 2.8 Gabor Filters

In order to estimate the potential confounds introduced by presenting only a limited sample of stimuli we wanted to see what topology is generated by a simple gabor filter model presented with static gratings Jones and Palmer (1987). The Gabor filter is defined by the equation:

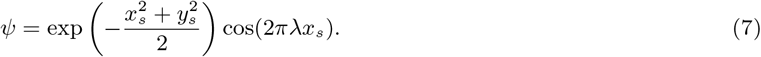

Where *x*_*s*_ = *x* cos *θ* + *y* sin *θ, y*_*s*_ = −*x* sin *θ* + *y* cos *θ, λ* is the frequency and *θ* is the orientation of the filter. To calculate the homology of the responses, we presented gratings with an increasing amount of orientations. Each grating was pointwise-multiplied with each filter, all pixels were then summed and a ReLU function *f* (*x*) = max(0, *x*) was applied to the output. The preferred frequency of the filters was sampled from an exponential distribution with a rate parameter of 0.69, the preferred orientation - from a uniform distribution between 0 and 2*π* and the output of each grating was scaled by a value from an exponential distribution with a rate parameter = 20 (this is done to resemble the different firing rate properties of neurons).

## 3. Results

### 3.1 When do standard methods fail?

Having created high dimensional non-linear manifolds with a known topology, we can start to evaluate the performance of the *persistent homology* algorithm using different metrics. Figure 1 shows a 2 dimensional PCA embedding of a 9 dimensional manifold, in which each principal component contributes equally to the explained variance. In this situation one would expect to find a single circle, but the Euclidean metric fails because it is a bad approximation of the true geodesic distance (panel C). As a result, the topological features which one can extract with homology will fail to correctly reflect the true structure of the manifold from which the point cloud was sampled. On the other hand using the numerical calculation of the geodesic correctly indicates the presence of a single 1 dimensional hole.

**Figure 1:**
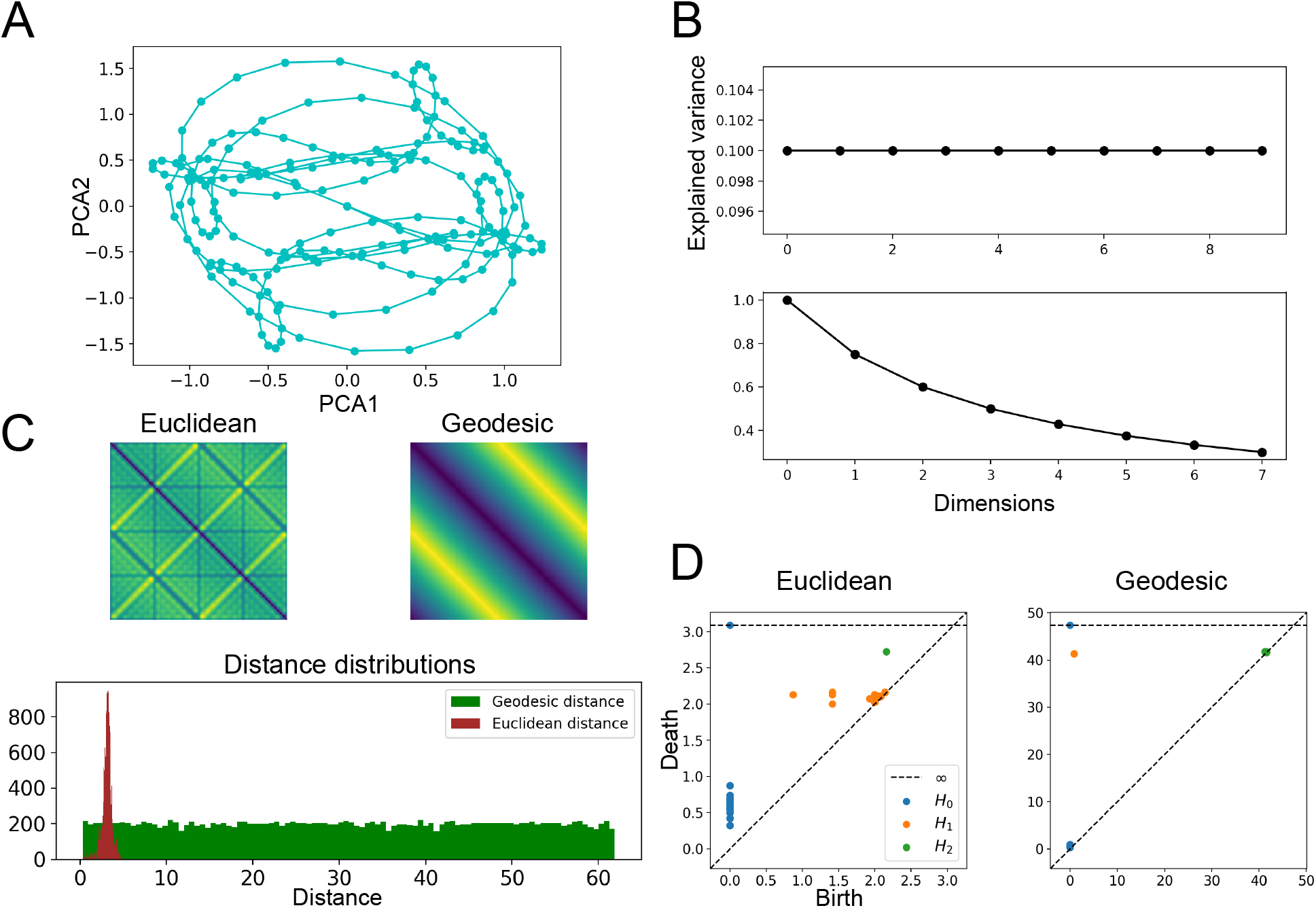
A) shows a PCA embedding of the high dimensional circle created by orthogonal basis functions. B) the top plot shows the explained variance ratio by each principal component within a 9 dimensional manifold while the bottom plot shows the explained variance by the first three principal components as extra dimensions are added, the first three components explain 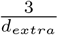 of the total variance. C) The top two plots show the distance matrices for both the Euclidean and the geodesic distance, while the bottom plot shows the distribution of distances for the two metrics. D) Shows persistence diagrams for both the Euclidean and Geodesic metrics.

Some other important properties which determine whether one should use a geodesic metric are the number of samples, curvature and sampling noise. As can be seen in figure 2, panel A, initially the difference between the metrics increases rapidly, but after a certain number of samples increasing them further leads to a similar performance of the *persistent homology* algorithm for both metrics. This shows that in the geodesic case, less samples are necessary to achieve correct identification of the ground truth topology. This plot also begs the question why even as the difference in the distance distribution increases, the difference in the obtained homology becomes smaller. The increase of the distance difference with the number of samples is expected, because denser sampling leads to a better approximation of the geodesic distance, which is equal or larger than the Euclidean distance between any two points. This fact implies that the errors due to the metric are problematic - only when they find shortcuts which qualitatively change the topological nature of the data.

**Figure 2:**
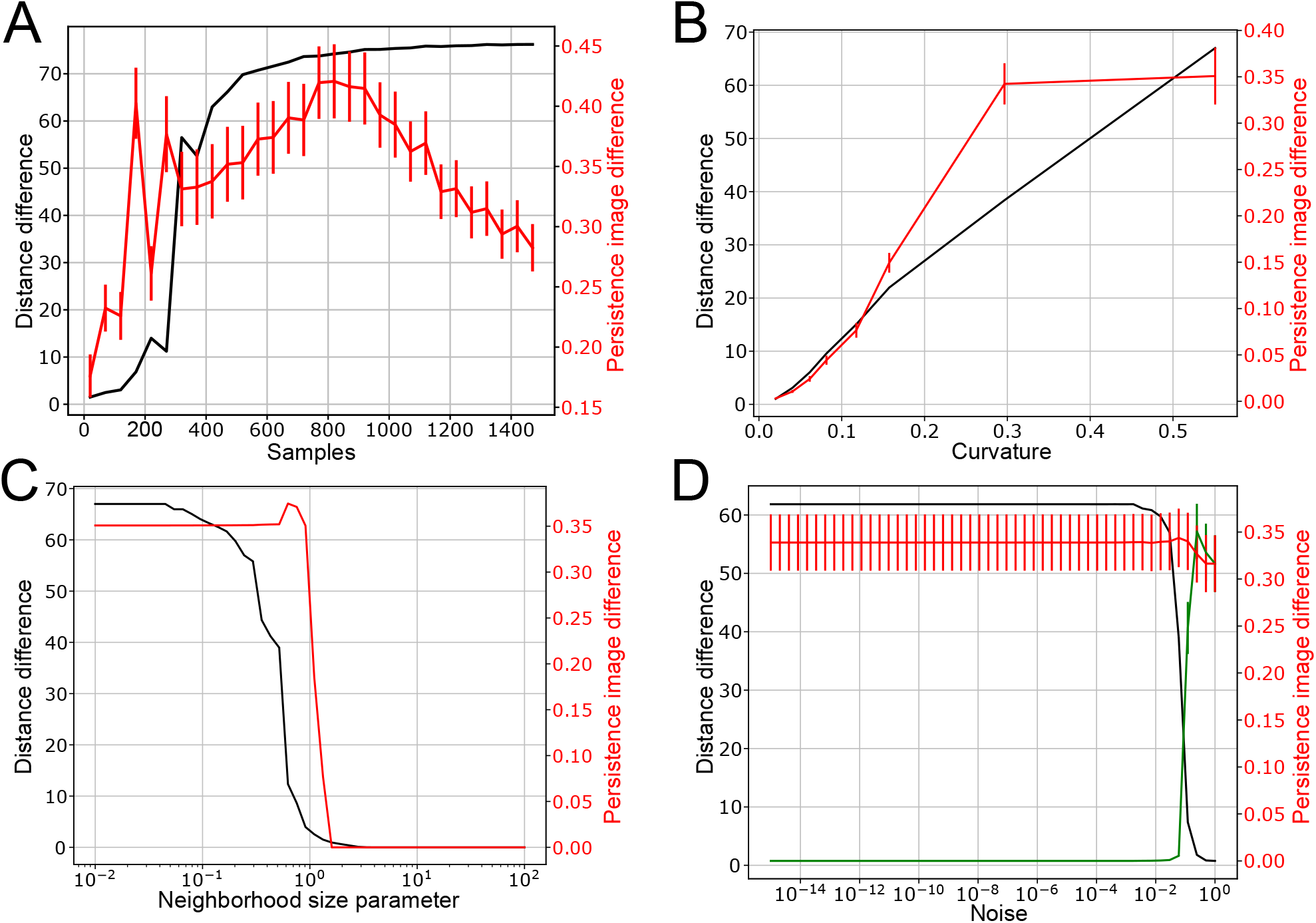
A) shows how the difference in the distance distributions (black line) and homology (red line) changes between the geodesic and Euclidean metrics as a function of the number of samples, the errobars reflect the SEM of the error distributions. B) Distance and homology difference as a function of curvature, defined as the average of the absolute value of the curvature at each point. C) Distance and homology difference as a function of the neighborhood parameter. D) Distance and homology difference as a function of the noise added to the manifold. The green curve shows the difference between persistence images obtained with the geodesic metric for zero noise and the geodesic metric for higher noise, the red curve shows the difference between the geodesic metric for zero noise and the Euclidean metric for higher noise values, as before the black curve shows the difference between the distance distributions of the noisy samples.

Of course, here we show how the topological structure which one can identify depends on the parameters for a single explicitly constructed manifold and generalising these results to any manifold is a difficult task. Nevertheless our results point to the fact that the geodesic distance outperforms the Euclidean distance when few samples are available and performs equally well for a large number of samples. A higher curvature implies that the geodesic distance will deviate more from the Euclidean distance. As a result, in panel B, we see that the difference between the metrics grows as a function of the curvature and the difference in the homology follows the same pattern. In panel C, we see that as the neighborhood parameter is increased the geodesic distance starts to converge to the Euclidean distance and the features identified through homology closely follow. In panel D, we see that the difference between the metrics is not affected by small amounts of noise, but for high amounts of noise the manifold starts looking like a random point cloud in Euclidean space. In that case the geodesic metric does not help, as the manifold structure is in essence Euclidean.

### 3.2 Extracting features from real data

We ran the *persistent homology* algorithm on data from 5 regions across 54 mice with both the geodesic and Euclidean metrics. We found several significant topological features with both methods. An example analysis of a region is shown in figure 3. As one can see in panel B, according to our 80% explained variance criterion, quite a large portion of the manifolds that we find in the data are also high dimensional and nonlinear, which justifies using the geodesic approach.

**Figure 3:**
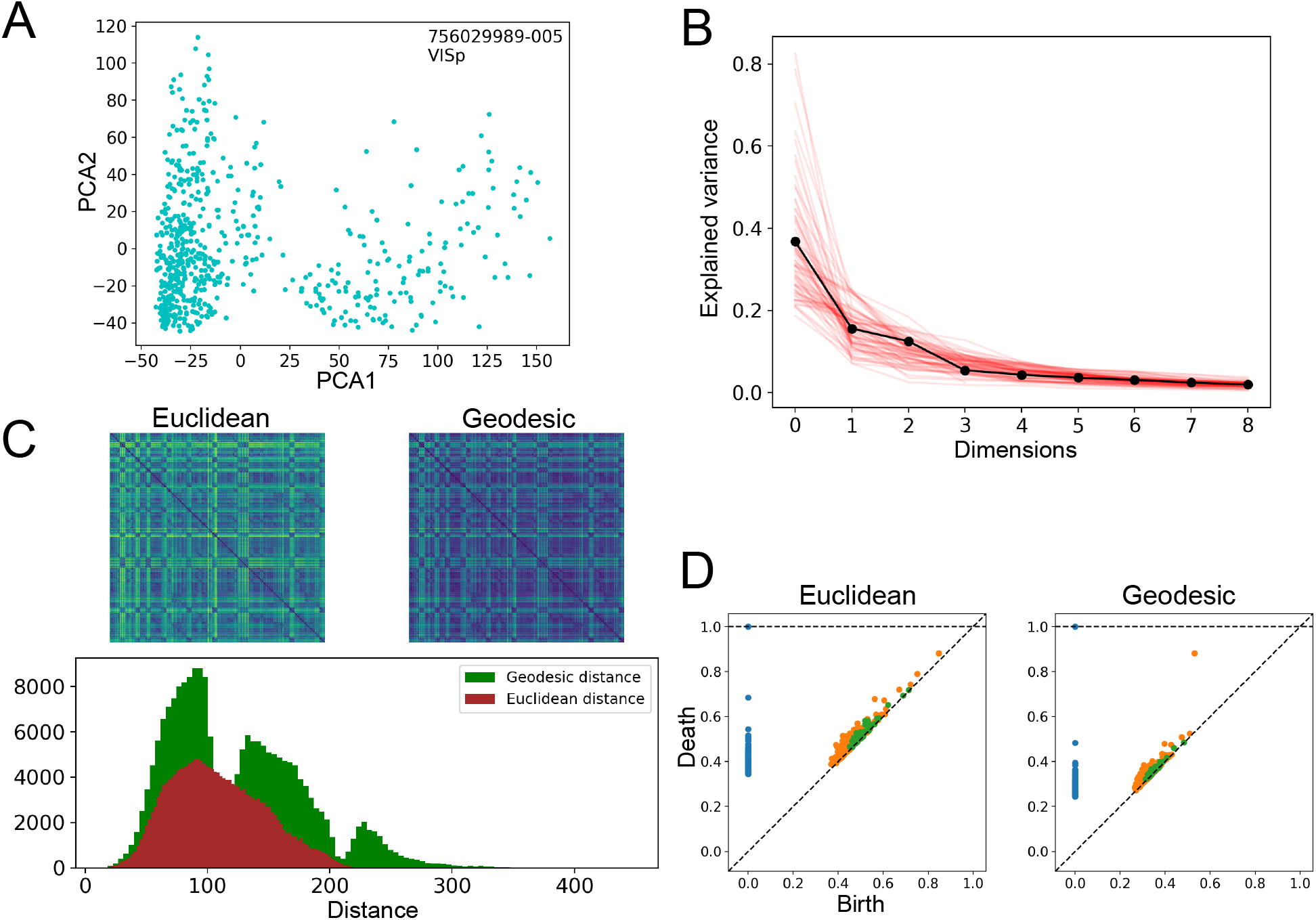
A) shows a PCA embedding of one experimental dataset with a significant feature detected only by the geodesic approach. B) shows the explained variance ratio for the first 9 principal components. C) The top two plots show the distance matrices for both the Euclidean and the geodesic distance, while the bottom plot shows the distribution of distances under the two metrics. D) Shows the persistence diagrams for both the Euclidean and Geodesic metrics.

The comparison of the two metrics shows that when using the geodesic distance one finds larger features and that, depending on the neighborhood parameter, one can also find more statistically significant features. Interestingly, there are also features which are visible only when using the Euclidean distance, which means one should try several neighborhood parameters when analysing a specific neural manifold. These observations are documented in figure 4.

**Figure 4:**
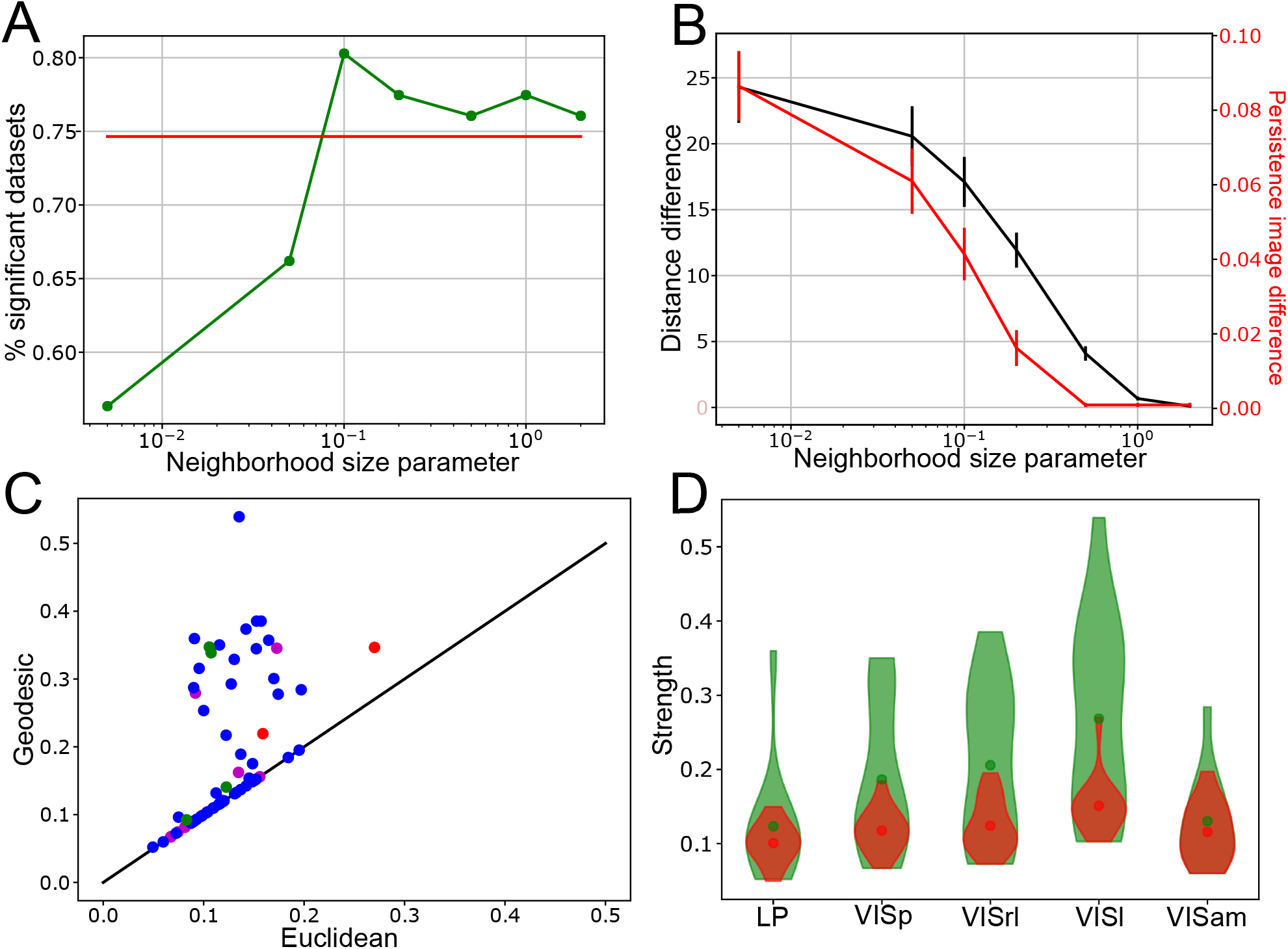
A) shows the fraction of significant 1 dimensional holes (across 71 datasets) for both the Euclidean (red) and Geodesic (green) metrics for different values of the neighborhood parameter. B) Shows the average difference between the distance matrices and perisistence images for all datasets as a function of the neighborhood parameter. C) Shows the strength and significance of the strongest feature in each dataset under both metrics, with a neighborhood parameter value of 0.1. The significance is color coded - significance under both metrics (blue), significance only under the geodesic metric (green), significance only under the Euclidean metric (red) and no significance (magenta) D) Violin plots for the strength of the strongest topological feature in each region, like above - green is used for the geodesic distribution and red - for the Euclidean.

### 3.3 Topological features of the visual system

Fixing the neighborhood parameter at 0.1, chosen from the peak of figure 4A, we show how manifolds with differing topology are distributed across visual regions. In figure 5C we see that in each region there are several cases in which the manifold has a complicated topology, that is not easily classifiable into a simple manifold like connected space, the circle, the sphere or the torus. These more complicated manifolds can be multiple circles or spheres as well as combinations of the two. While there are clear differences between the manifolds that appear, we weren’t able to find any significant differences between the features in any of the regions. Of course it is hard to establish statistical significance for such a small amount of samples, which might explain why we don’t detect more regional differences. On the other hand we found strong effect sizes between the bootstrap distributions in most comparisons (figure 5B), which leads us to believe that given a larger sample of subjects one would be able to find significant differences between the regions.

**Figure 5:**
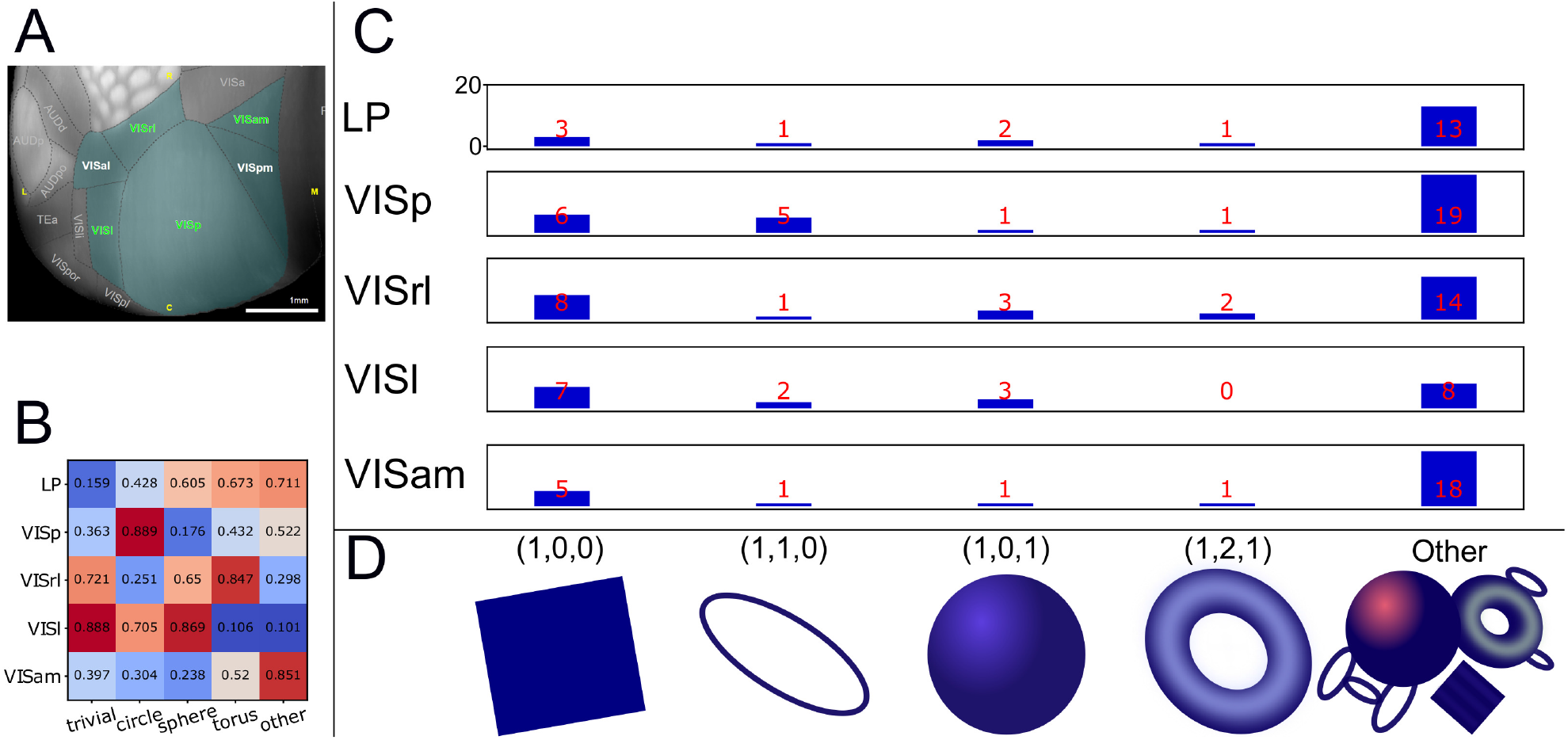
A) A cortical flatmap of the regions included in our analysis (names in green). In addition we analyse one thalamic region (LP) which is not shown here. Image credit: Allen Institute. B) P values of the difference between the topological features found across the different regions C) Shows the number of manifolds with a particular topology throughout the datasets. D) The manifolds corresponding to the bars above them. The numbers in the brackets show the Betti numbers, which characterize the homology of each manifold. The “other” category contains several different manifolds, including tuples of circles and spheres, as well as more complex combinations of the two.

We also looked at the manifolds generated by drifting gratings with a fixed temporal frequency. In these conditions the only relevant stimulus parameter is the orientation of the grating and if a neural network codes for the orientation, one would expect to find circle-like manifolds. The manifolds we found when including data for all temporal frequencies are shown in figure 6. Panel C shows the average change in the amount of manifolds found between the “all temporal frequencies” condition and the fixed frequency conditions. It is clear that when the temporal frequency is fixed, the explored manifold has a circular structure more often and in turn the amount of manifolds with complicated topology decreases. One explanation for this results is that neural activity is exploring a simpler submanifold on which orientation is encoded. In addition, in the frequency fixed case, we see that there is an average increase in the amount of circles and a decrease in the amount of “other” more complicated manifolds. In figure 7, we show for which stimulus/region combinations this change in topological features is significant.

**Figure 6:**
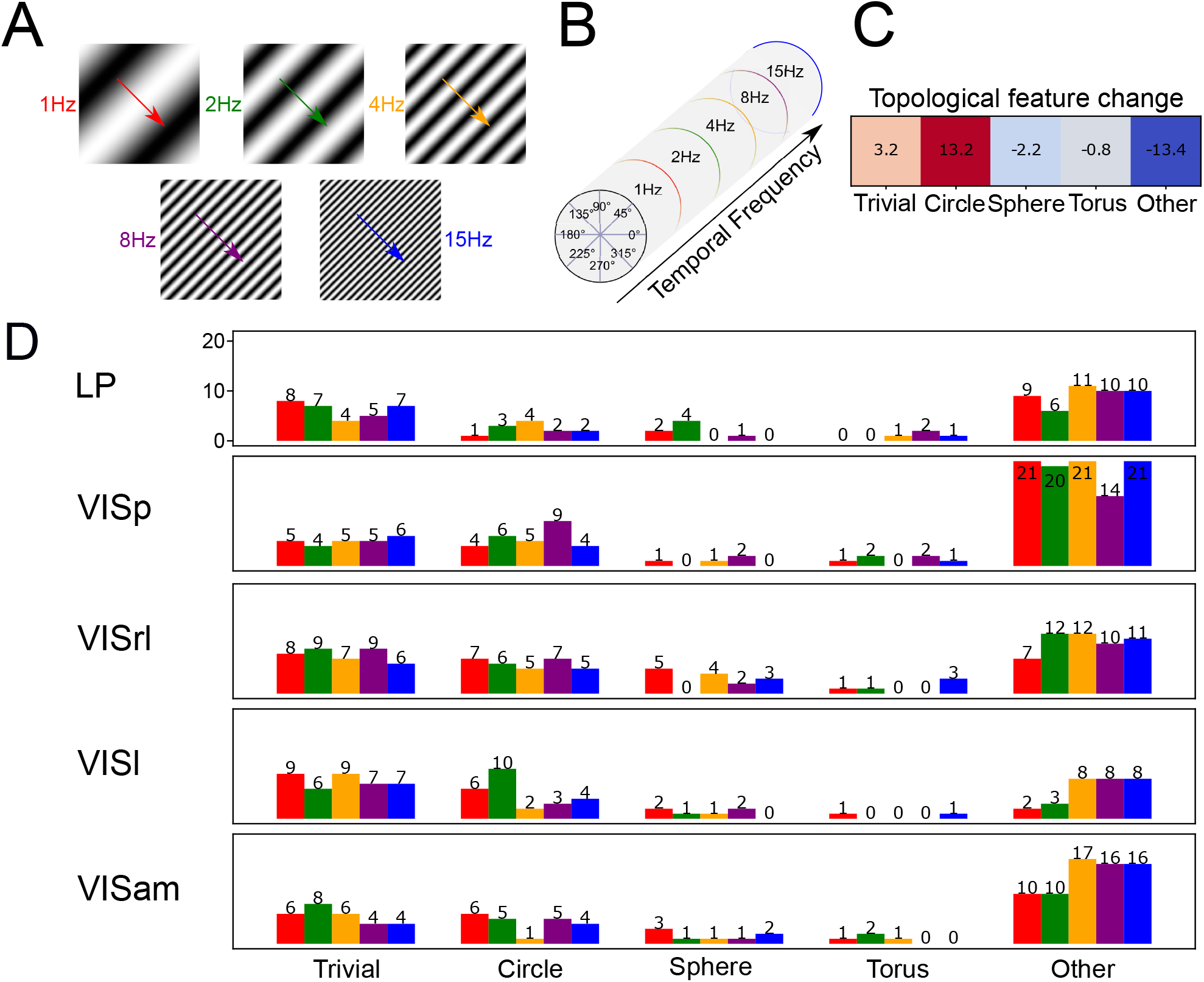
A) Examples of the drifting grating stimuli. The increasing temporal frequency is shown as spatial frequency (which is constant at 0.004 cpd). B) A diagram showing the different stimulus parameters. The cylinder shape comes from the fact that one variable is periodic (orientation) while the other is not (temporal frequency). C) Shows the mean change between the topology of the manifolds under all stimuli (each column of bars in figure 5 C) and the stimuli with fixed temporal frequency (all 5 bars in a column). D) The manifolds corresponding to the bars above them, color coded for the temporal frequency at which the stimuli were fixed.

**Figure 7:**
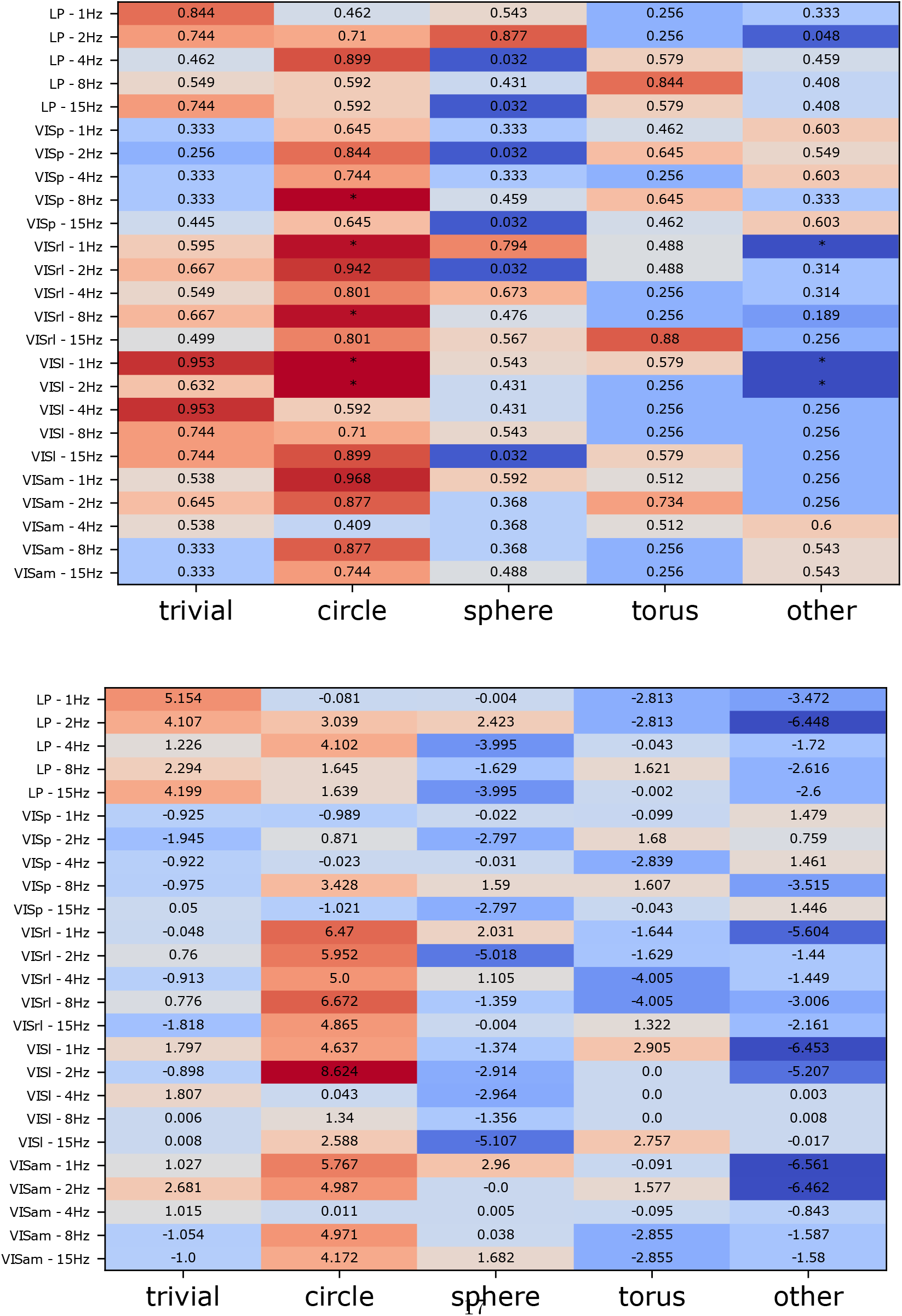
False discovery rate corrected p values (left - stars denote significance for p*<*0.025) and Cohen d based effect sizes (right) of the comparisons between the distributions of manifolds under all stimuli and those at specific fixed temporal frequencies.

## 4 Discussion

We have shown that there are cases in which treating neural activity as a simple low dimensional manifold will miss important and interesting structure that is present in the manifold. Furthermore when applying *persistent homology*, the Euclidean distance creates artificial shortcuts which are not reflective of the paths on the manifold that connect two points. In the neuroscience literature there is a view that the topological structure of a neural manifold reflects the coding or dynamic properties of a neural population (Curto (2017)). While we agree with this notion, finding such manifolds in practice comes with serious challenges. The *persistent homology* approach comes with the limitation that even relatively simple manifolds like circles, spheres and tori can be deformed in complicated ways without changing their topological properties. Furthermore we usually sample an insufficient number of points from such neural manifolds. As a result one should expect to miss true positive or find false positive results due to the choice of an inappropriate metric. The geodesic metric is the distance metric which makes the weakest assumptions about the manifold and since one can vary the neighborhood parameter, it is also a more general metric than the Euclidean one (it even becomes equivalent to the Euclidean metric for very large neighborhood values). However this generality doesn’t come without a price.

The fundamental limitation of our approach is that one has to choose a value for the neighborhood parameter and this choice reflects the researchers’ belief of the scale at which the manifold is well defined. A small value implies the belief that the point cloud is sampled from a tight distribution, while a larger value implies a more smeared out underlying distribution. If the point cloud one is looking to analyse is not too big, one can use a continuous range of neighborhood parameters and see if there is a stable range of parameter values which produce interesting structure. Another approach is to say that we want to allow for the broadest distribution while avoiding the problem of artificial shortcuts. There is some work related to the Isomap algorithm (Tenenbaum et al. (2000)), which proposes to exclude outlier points (Choi and Choi (2007)) in order to avoid topological instability, due to such shortcuts.

The neural data that we analysed showed that, at the very least in the mouse visual system, the assumption of neural data being distributed on simple low dimensional manifolds is rarely true when a large amount of neurons are included. Using our geodesic approach to *persistent homology* we find that the manifolds generated by the visual system in response to drifting gratings with varying temporal frequency, can range from simple connected spaces to ones with a very complicated topology. Following the results in (Stringer et al. (2019)), where it is shown that in order to preserve smoothness, the decay of the eigenspectrum of a neural population has to decay faster than 1 + 2*/d* (where d is the number of stimulus parameters), we would expect that presenting complex stimuli like natural scenes will lead to the predominant identification of even more high dimensional and topologically complicated manifolds.

Singh et al. (2008) performed a similar analysis to ours, except for the fact that their analysis was done in monkeys responding to natural movies and involved a much smaller number of recorded cells covering the same patch of visual space. Our results find that there are much more complicated manifolds to be found when the number of recorded neurons increases and they cover a large portion of visual space. This result makes sense, even though cells in the visual system have strong receptive fields, it has also been found that around 34% of cells don’t have any tuning properties (de Vries et al. (2020)). One could still expect that the responsiveness of visual cells is dependent on internal variables which lead to a manifold topology dependent on the combination of the topological structure of incoming stimuli and that of internal variables, which is unknown. In addition we see that there are several cases (8% of all datasets) in which the manifold topology is the same as that of a sphere, which is consistent with the proposed model in (Bressloff and Cowan (2003)). We achieve this results with a fixed spatial frequency. According to the energy model of V1 Mante and Carandini (2005) the same topology should be achievable with varying temporal frequency. Under that interpretation of visual activity one would expect to find circles when the temporal frequency is fixed as then the only parameter of importance is orientation. Nevertheless, the fact that such manifolds were not found consistently across these datasets could be explained by the fact that analysing a significantly higher number of cells leads to the inclusion of neurons that code for other properties, which distort the spherical manifold.

Another explanation for the lack of clear manifold structure is the low density of stimulus sampling, by which we mean the amount of stimuli presented. Only 4 orientations (8 directions) were presented, which is equivalent to sampling only 4 points from the stimulus manifold. While 4 points is the minimum amout of data required to identify a hole in a point cloud and these points are sampled repeatedly across trials, the stimulus space is still sampled only very sparsely. Even if we interpret higher order visual processing as simply applying Gabor filters to stimuli, 4 orientations would not be enough to consistently find circles, which are easily found with higher stimulus sampling (figure 8).

**Figure 8:**
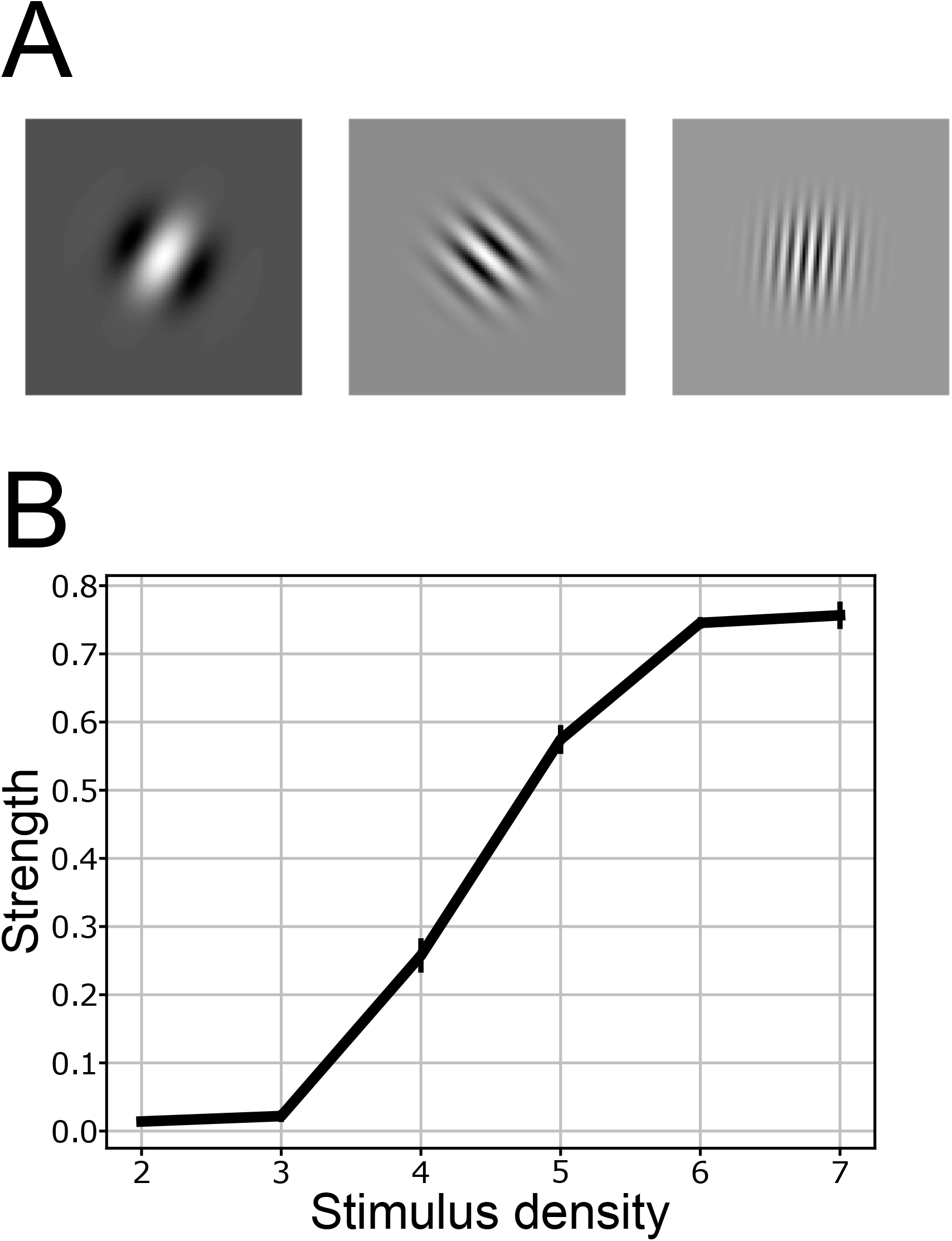
A) Example Gabor filters which are presented with static gratings. B) Stimulus density vs the persistence of the identified circular feature, where stimulus density (*s*) is defined as 2^*s*^, which is the amount of presented orientations to the Gabor filters.

These observations points to the conclusion that neural activity in the visual system explores a very complicated manifold and that by presenting a small subset of stimuli, one can only find the topology of a submanifold. In the future it would be interesting to see how the topology of these manifolds changes if one increases the number of presented stimuli. This can be done using models, that have been shown to perform similarly to the currently available data and allow for the presentation of custom stimuli (Billeh et al. (2020)). Our results on the analysis of neural manifolds in response to fixed temporal frequency stimuli support the hypothesis that, when presented with a subset of stimuli, neural networks explore submanifolds on which particular stimulus parameters are being encoded. In theory the stimulus manifold of orientation and temporal frequency should have the topology of a cylinder which has the same homology groups as a circle. However the fact that we see an increase in the number of circles as a result of fixing the temporal frequency, implies that the visual system does not perform a continuous transformation of the original cylinder stimulus manifold. Lastly, despite the fact that different regions have been shown to have different orientation and temporal frequency tuning (Andermann et al. (2011), Marshel et al. (2011)), we were not able to find significant differences in the topology of their regional activity. This could either mean that the topology of the regions is not different despite the difference in their tuning properties or that more data is needed. In any case the high effect size values Cohen (2013) point towards the possibility that a more targeted analysis can find differences in the topology between regions. Increasing the sampling density of temporal frequency and orientation in the stimulus set, could help bring more clarity to these questions.

In summary, we propose that if one wants to find more intricately deformed non-linear high-dimensional manifolds with *persistent homology*, the geodesic metric might be a good tool to use, while the Euclidean metric might lead to inaccurate results. Furthermore we show how our approach depends on a number of fundamental properties of the point cloud on the manifold from which one samples, like the sampling density, curvature, neighborhood size and sampling noise. Lastly we confirm the usefulness of the geodesic metric by finding significant topological features in neural data, which are missed by the Euclidean metric. These features reveal interesting information about how stimulus manifolds are mapped to neural activity in the mouse visual system. Further exploration of this question can significantly help our understanding of stimulus representations in the brain.

